# Exploring contemporary and historic effective population sizes in Atlantic beluga whale populations

**DOI:** 10.1101/2025.09.19.677402

**Authors:** Claudio Müller, Evelien de Greef, Cortney A. Watt, Steven H. Ferguson, Veronique Lesage, Geneviève J. Parent, Colin J. Garroway

## Abstract

Ongoing global environmental changes will require animals to adapt to new conditions, as their necessary resources for survival may shift or decline. Understanding population dynamics in wildlife informs conservation strategies by providing context for genetic diversity and current population abundances. Here, we use whole genome data to reconstruct past demographic trends and estimate contemporary effective population sizes in the beluga whale (*Delphinapterus leucas*), an Arctic endemic species with significant cultural importance, that is an integral part of the marine ecosystem along the Canadian coast. In line with previous work, we identified six genetic populations: 1) St. Lawrence Estuary, 2) Eastern High Arctic, 3) James Bay, 4) Cumberland Sound, 5) Hudson Bay-Strait Complex and 6) Little and Great Whale Rivers. Demographic history in the deep (up to 1,000,000 years) past revealed all but the Hudson Bay-Strait Complex population declined in effective population sizes following the last glacial period. Within the last several centuries, the Eastern High Arctic and Hudson Bay-Strait Complex populations have shown evidence for an increase in effective population size and have contemporary estimates greater than 2,000. The other four populations have contemporary estimates below 400. The very low effective population size estimates in Cumberland Sound, Little and Great Whale Rivers and James Bay, as well as the high inbreeding in the St. Lawrence Estuary indicate that these populations are in need of protective measures for long-term sustainability. These results have advanced our understanding of the dynamics of beluga genetics across eastern Canada and provide valuable metrics for conservation strategies.

## Introduction

Climate warming has altered species’ abundances and distributions globally (Díaz et al., 2019; Pörtner et al., 2023; Scheffers et al., 2016). Some species have responded to warming through combinations of tracking climatic niches, phenotypic plasticity, and adaptive evolution (Chen et al., 2011; Hoffmann and Sgrò, 2011; Urban et al., 2024). In contrast, many other species face habitat losses with associated population declines and increased extinction risk (Maclean and Wilson, 2011; Urban et al., 2024; Urban, 2015). In general, this reorganization of biodiversity is substantially altering the ecosystems that sustain wild species and people (Díaz et al., 2018). In the Arctic, the rate of warming is approximately four times the global mean (Rantanen et al., 2022). The threats to Arctic ecosystems associated with rapid warming are compounded by the high degree of specialization of Arctic species on disappearing features of Arctic environments (e.g., sea ice) and increases in industrial development due to growing access to the region (Laidre et al., 2015). Information about the current ecological and evolutionary status of wild populations is necessary for assessing risks posed by warming and understanding the consequences of future change, especially in highly sensitive ecosystems.

A central goal of conservation and management is to maintain the adaptive capacity of wild populations (Armitage and Plummer, 2010; Mcleod et al., 2016). Patterns of genome-wide diversity through space and the rates at which diversity is lost across populations underlie the microevolutionary processes that contribute to population mean fitness and the resilience of a population to environmental changes (Lande and Shannon, 1996). All populations lose diversity due to random genetic drift (Li and Graur, 1991). The evolutionary concept of *effective population size* (*N*_e_) provides an estimate of the strength of genetic drift and the consequent rate of loss of genetic diversity within a population (Wang et al., 2016; Waples, 2025, 2022). Additionally, the strength of genetic drift is inversely proportional to the efficiency of natural selection (Charlesworth, 2009; Kimura, 1983; Ohta, 1992). As effective population sizes decline, the random process of genetic drift can increasingly overwhelm the deterministic process of natural selection. Knowledge of the past and present trends in effective population sizes is thus important for prioritizing conservation and management decisions in light of future environmental change.

Beluga whales (*Delphinapterus leucas*) are a vital part of the Arctic food web (Heide-Jørgensen and Teilmann, 1994; Kelley et al., 2010; Seaman et al., 1982) and have been found to be an indicator species for environmental changes (Wagemann et al., 1995). Belugas also hold cultural importance for northern Indigenous communities, who rely on them for food, materials, and cultural practices (Hoover et al., 2013; Tyrrell, 2008). In some regions, they are also important for tourism (Dressler et al., 2001), which supports local economies (e.g., Churchill; The Churchill Beluga Whale Tour Operators Association et al., 2015). However, recent centuries have seen a decrease in the number of belugas in some regions, driven in most cases by commercial whaling (Harwood et al., 2014; Mitchell and Reeves, 1981; Reeves and Mitchell, 1984; Rugh et al., 2010; Tinker et al., 2024; Watt et al., 2021). While Indigenous communities have hunted whales for millennia, large-scale commercial whaling by Western settlers began in eastern Canadian waters in the early 16^th^ century (Loewen, 2009; Tuck and Grenier, 1981). Commercial whaling peaked with respect to beluga harvesting in the early or late 19^th^ century (Bockstoce, 1986; Reeves and Mitchell, 1984) and continued until bans in the Arctic in 1972 (DFO, 1981; International Whaling Commission, 1982) and the St. Lawrence Estuary in 1979 (Reeves and Mitchell, 1984). An example demonstrating the scale of harvest is provided by Stewart (2018), who estimated that a minimum of 14,079 beluga whales were commercially harvested from Cumberland Sound alone between 1860 and 1966. These numbers are likely underestimates, as they do not account for hunted whales that were not recovered. Indigenous harvests during this time are not well-documented; however, commercial harvest in some years was more than 15 times larger than current subsistence catches (Stewart, 2018).

The genetic consequences of commercial whaling have been observed in bowhead whales (*Balaena mysticetus*) (De Greef et al., 2024), also an endemic Arctic species, demonstrating that overexploitation can have negative effects on genetic diversity. Today, only Indigenous communities are permitted to hunt belugas (Meehan et al., 2017) but the small size of some of the populations may lead to unsustainable harvest (Van de Walle et al., 2025). Additionally, these animals face increasing threats due to climate change. Heavily reliant on sea ice to protect them from orcas (*Orcinus orca*), belugas experience an increased threat of predation as sea ice melts due to rising temperatures (Breed et al., 2017; Ferguson et al., 2010; Higdon et al., 2012; Stafford, 2019). Additionally, changes in beluga aggregation behaviours have been directly linked to the increase in sea surface temperature, leading to reduced presence of whales in warming waters (Rivas et al., 2024). In the St. Lawrence Estuary, the southernmost limit of the beluga worldwide distribution, calf mortality has been related to the increase in water temperature and reduction in sea ice indices (Williams et al., 2021). Predictive models indicate a northward shift in beluga habitat, and a reduction in the range of Arctic whales in general (Chambault et al., 2022).

Beluga whales in Canada have been divided into eight stocks for management purposes. These stocks are based on summering aggregations due in large part to individual beluga whales’ high site fidelity to summer locations across years (Finley, 1982). These summer areas include shallow coastal estuaries and bays that may offer abundant food (Seaman et al., 1982), warmer waters that facilitate skin moulting (St. Aubin et al., 2011), and protection from predators during birthing and care of calves (Fraker et al., 1979; Sergeant, 2011; Sergeant and Brodie, 2011). Seven beluga stocks occur in eastern Canada, and one resides in the western Canadian Arctic in the Eastern Beaufort Sea (COSEWIC, 2020) (Figure 1).

**Figure 1:**
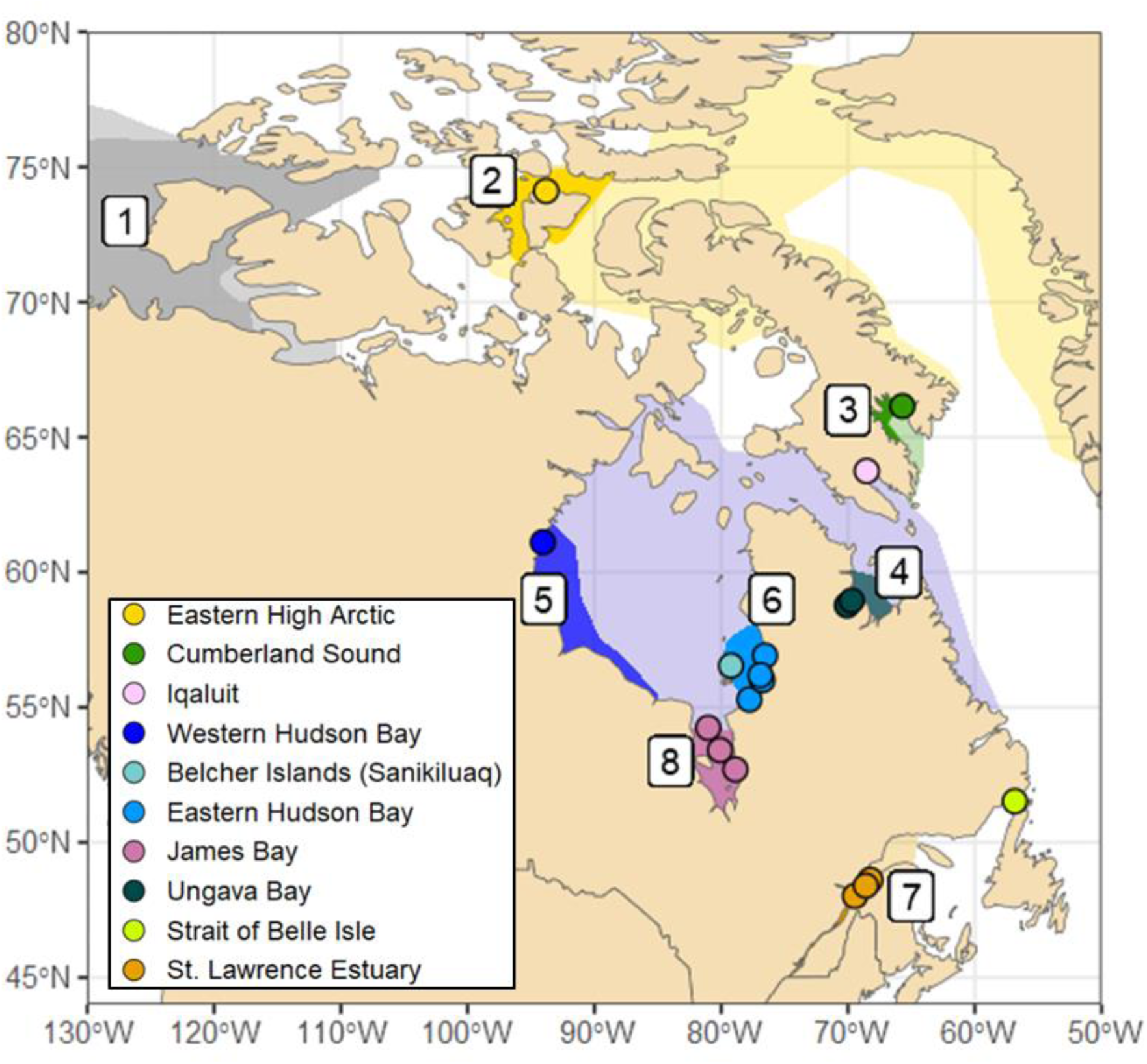
Canadian beluga stocks and range (COSEWIC, 2020). Dark colored areas show summer habitats; lightly colored areas show seasonal migration range. Designated stocks are: (1) Eastern Beaufort Sea, (2) Eastern High Arctic, (3) Cumberland Sound, (4) Ungava Bay, (5) Western Hudson Bay, (6) Eastern Hudson Bay, (7) St. Lawrence Estuary and (8) James Bay. The dots show the sampling locations: Eastern High Arctic (n=15), Cumberland Sound (n=13), Iqaluit (n=10), Western Hudson Bay (n=15), Sanikiluaq (n=15), Eastern Hudson Bay (n=29), James Bay (n=16), Ungava Bay (n=3), Strait of Belle Isle (n=1) and St. Lawrence Estuary (n=29).

The most threatened of these stocks are the Eastern Hudson Bay, Cumberland Sound, Ungava Bay, and St. Lawrence Estuary stocks (COSEWIC, 2020; Lesage et al., 2024). The latter three stocks have been listed as endangered by the Committee on the Status of Wildlife in Canada (COSEWIC) and are now under special management. Several recent aerial surveys of the Ungava Bay stock found no or a few tens of beluga whales in the region, questioning the persistence of this stock (Sauvé et al., 2024). It is unclear whether individuals sampled in that region today represent a small number of remaining individuals from the original stock or whether they now consist of individuals that have moved into the region from Hudson Bay and use the area for migration (COSEWIC, 2020). Hunting quotas for Eastern Hudson Bay and Cumberland Sound beluga are in place to manage sustainability and reduce further decline (DFO, 2024; Stewart and Lockhart, 2005). The St. Lawrence Estuary stock is of particular interest because they are geographically disjunct and occur south of the Arctic (Brown Gladden et al., 1997), having diverged from other beluga populations after the end of the last ice age (Harington, 1977). The St. Lawrence Estuary stock was subject to substantial hunting and eradication programs following the European colonization of the region (Reeves and Mitchell, 1984; Vladykov, 1944). Belugas were hunted for their fat, skin, and oil, and in the St. Lawrence Estuary, they were also killed because they were seen as competing with fishermen for fish (Pippard and Malcolm, 1978). Due to large declines, the hunting of belugas in the St. Lawrence Estuary has been forbidden entirely since 1979 (Kingsley, 2002; Reeves and Mitchell, 1984). Despite evidence of demographic stabilization in some of these endangered populations, there is little indication of recovery, and projections point toward the possibility of further declines (Tinker et al., 2024; Van de Walle et al., 2025; Watt et al., 2021).

Beluga population trends are difficult to assess directly due to their remote location, especially in the Arctic, the considerable amount of time belugas spend underwater and under dense sea ice, and their clumped distribution, which often results in large variances around abundance estimates (St-Pierre et al., 2024). Additionally, most beluga whale populations migrate during the fall and spring (Figure 1). As belugas mate during late winter and early spring while travelling to their summer habitat (O’Corry-Crowe et al., 2018, 1997), gene flow may occur between stocks even though their summer habitats are geographically isolated. Studies based on mitochondrial DNA and microsatellites have provided evidence for genetic divergence between the stocks, maternal site fidelity, and natal philopatry (Brown Gladden et al., 1997; Gladden et al., 1999; Skovrind et al., 2021; Turgeon et al., 2012). A recent genome-scale study identified six genetic beluga whale populations in eastern Canada, occurring in the Eastern High Arctic, Cumberland Sound, Hudson Bay-Strait Complex, Little and Great Whale Rivers, James Bay, and St. Lawrence Estuary (Montana et al., 2024). Contemporary and recent changes in effective population size in these populations have not yet been explored.

To better understand threats to genetic diversity and the adaptive capacity of beluga whales, we used whole genome data from beluga sampled across eastern Canada to explore their demographic histories and contemporary effective population sizes. After determining the underlying population structure of our samples, we reconstructed changes in effective population size (*N_e_*) for each population in the deep (up to 1,000,000 years) and recent (up to 3,000 years) past. Finally, we estimated the contemporary *N*_e_ within the context of demographic histories and the future capacity for adaptive change.

## Methods

### Tissue collection, extraction & sequencing

In this study we used samples from 15 locations, representing seven of the eight Canadian beluga stocks: St. Lawrence Estuary, Ungava Bay, Eastern Hudson Bay, Western Hudson Bay, James Bay, Cumberland Sound and Eastern High Arctic (Figure 1). Skin tissues were collected from beluga whales harvested by Indigenous subsistence hunters between 1990 and 2020 (n=116), or from beluga carcasses found on the beach in the St. Lawrence Estuary or Strait of Belle Isle (n = 30). The DNA from 78 individuals (sampled from Eastern High Arctic, Cumberland Sound, Iqaluit, Western Hudson Bay, Sanikiluaq, and St. Lawrence Estuary) were extracted using the Qiagen DNeasy Blood & Tissue kit using the tissue extraction protocol. They were sequenced with Illumina NovaSeq to obtain whole genome paired-end reads with a target coverage of 10x, available at BioProject ID PRJNA1182041 (De Greef et al., 2025). The data for 68 individuals (sampled from Eastern Hudson Bay, James Bay, Ungava Bay, and named locations within St. Lawrence Estuary (Table S1)) were obtained from BioProject ID PRJNA984210 (Montana et al., 2024).

### Alignment, variant calling & baseline SNP filtering

We trimmed the raw sequences using *Trimmomatic* v0.36 (Bolger et al., 2014) and aligned the sequences using *Bowtie2* v2.5.2 (Langmead and Salzberg, 2012) to the reference genome assembly GCA_029941455.3 (Bringloe and Parent, 2023). Duplicate reads were removed and read group information was added using *Picard* v2.20.6 (Broad Institute, 2019). Given our coverage variation, we used *GATKv4.1.2* (McKenna et al., 2010) to down-sample the high coverage samples to a target modal coverage of 6x. Genomic variants were called using *Platypus* v0.8.1 (Rimmer et al., 2014). We then used *vcftools* v1.16 (Danecek et al., 2011) and *bcftools* v1.19 (Danecek et al., 2021) to filter out indels, sites without the PASS flag, low quality sites (QUAL<50), sites with missingness > 0.25, non-biallelic sites and scaffolds shorter than 100kbp as well as scaffolds aligned with the sex chromosome to obtain only high-quality autosomal Single Nucleotide Polymorphisms (SNPs).

Then, we used *PLINK* v1.9 (Purcell et al., 2007) to assess kinship and removed the individual with the higher rate of missing data from pairs with pi-hat > 0.4 to remove bias on analyses from first-degree relatives. We also removed samples with more than 30% overall missing data. After these filtering steps, 140 samples remained in our final data set (Table S2).

### Population structure & runs of homozygosity

To estimate genetic population structure, we used SNPs which are not under putative selection and not strongly linked. We filtered sites in Hardy-Weinberg disequilibrium (heterozygous frequency threshold > 0.6), with low minor allele frequency (< 0.05), and then pruned the data set using a minimum distance of 1,000 bp between sites (Table S3) using *vcftools* v1.16 (Danecek et al., 2011).

Using the remaining SNPs (1,081,765), we conducted PCAs using the r-package *adegenet* v2.1.10 (Jombart, 2008; Jombart and Ahmed, 2011) to examine population structure. We also estimated population structure with admixture analyses with K values 2 through 7, representing number of ancestral populations, using the snmf function from the r-package *LEA* v3.3.2 (Frichot and François, 2015). We used *vcftools* v1.16 (Danecek et al., 2011) to estimate pairwise genetic differentiation (*F*_ST_) between the identified genetic populations.

Finally, we estimated runs of homozygosity (ROH) as a measure of inbreeding. The greater the quantity of long homozygous sections in the genome of diploid individuals, the more likely we expect there to be a close relationship between the ancestors that have contributed to these regions in the chromosome. Therefore, a greater proportion of ROHs is indicative of higher degree of inbreeding (Ceballos et al., 2018). To estimate ROHs, we used *PLINK* (Purcell et al., 2007) using a sliding window size of 300kb, with a minimum of 50 SNPs at a minimum density of 1 SNP/50kb defining a ROH, allowing up to 3 heterozygote sites per window. A gap of 1000kb between two SNPs was required for them to be considered in two different ROHs.

### Demographic history & contemporary effective population size

For analyses of *N*_e_, we further filtered SNPs to exclude unplaced scaffolds that were not assigned to a chromosome in the reference genome and excluded sites out of Hardy-Weinberg equilibrium (heterozygous frequency threshold > 0.6; Table S4).

To reconstruct changes in *N*_e_ over time, we examined models estimating historic *N*_e_ in deep time (over thousands of generations ago), as well as in recent history (within the last 150 generations). For estimating demographic history in deep time, we used the coalescent theory-based method *SMC++* (Terhorst et al., 2017). This method allows deep-time demographic history reconstruction using low coverage, unphased sequences. To distinguish between long runs of homozygosity and missing data, we used *bedops* (Neph et al., 2012) to create a masked file to include in the *SMC++* model. We ran *SMC++* for each genetically distinct population. We randomly chose an individual from each population to represent distinguished lineages (Table S1). For *SMC++*, we set the regularization penalty to 4.0, the non-segregating site cut-off to 100,000, used a thinning of 2,000 and timepoints from 10 to 1,000,000 generations. The mutation rate per generation for belugas was set at 1.65x10^-8^ (Westbury et al., 2019). The resulting generation numbers were multiplied by a generation time of 28 years/generation (Lowry et al., 2019) to convert generations into years.

To estimate *N*_e_ over the last 150 generations, we used a linkage disequilibrium (LD) based method implemented in the software package *GONE* (Santiago et al., 2020). We used a maximum of 20,000 SNPs per chromosome, 400 bins and 2000 generations for which linkage data is obtained in bins. We ran 500 iterations per population (see Table S5 for complete parameter settings). Conversion to from generations to year was done again by multiplying by 28 years/generation.

Finally, to estimate contemporary *N*_e_, we used the LDNe estimator from the r-package *strataG* (Archer et al., 2017; Waples and Do, 2008). To prepare the SNP dataset for this analysis, we created a vcf-file per population that only contained the individuals from that population. We filtered for biallelic sites again and removed all sites with missing data. We used a minor allele frequency cut-off of 0.1 to avoid an overestimation of *N*_e_ resulting from rare alleles for populations with small sample sizes. We then randomly subsampled a set of 10K SNPs with *PLINK* (Purcell et al., 2007) 10 times to run multiple iterations with the LDNe estimator, then averaged resulting estimates of *N*_e_. Estimates of *N*_e_ are sensitive to underlying population structure, the presence of outlier individuals in a sample set, and inclusion of close kin (Macbeth et al., 2013; Novo et al., 2023; Wang, 2005). Thus, we estimated contemporary *N*_e_ for each genetically distinct population separately. Additionally, for populations containing strongly admixed individuals, we ran separate estimates including and excluding admixed individuals to better understand how these factors may impact results. We defined strongly “admixed” individuals as those that did not clearly fall into any cluster according to PC1 and PC2 in our PCA (Figure 1a). Given that all St. Lawrence Estuary belugas showed at least second-degree kinship with each other, we additionally estimated *N*_e_ for the St. Lawrence Estuary population with a subset of the 10 least related individuals.

While this approach allows us to test for bias introduced through admixed individuals, it is important to keep in mind that there are other sources of potential bias in our data set for *N_e_* estimates. As is common in wildlife applications, our samples include individuals from different, overlapping generations. Differences in sampling methods (e.g. subsistence harvest versus carcasses from shore) could also introduce biases if these sources tend to come from individuals genetically different from the general population. Finally, we have a range of sample sizes for each genetic population. While our use of whole genomes provides us with a large sample size in terms of alleles, *N_e_* estimates also benefit from a larger number of samples. These issues are well documented in (Waples, 2025).

## Results

### Population Structure

We identified six distinct genetic populations with both PCAs and admixture analyses (Figure 2). The St. Lawrence Estuary population was comprised of only belugas sampled from this region. The remaining individuals clustered as three large groups along PC2 (Figure 2a). The James Bay population consisted of all individuals from James Bay except those from Bear Island at the northern edge of the bay. This population also included five individuals from Kuujjuarapik in Eastern Hudson Bay and Sanikiluaq in the Belcher Islands (Figure S1). The Eastern High Arctic cluster consisted of all individuals sampled in the Eastern High Arctic and one individual sampled in Cumberland Sound. One remaining individual sampled in Cumberland Sound was admixed, showing shared ancestry with the Eastern High Arctic population and the third cluster, which comprised individuals from several stocks and sampling locations. Specifically, all individuals from Western Hudson Bay, Iqaluit and Ungava Bay, the one individual from Strait of Belle Isle in the eastern Gulf of St. Lawrence, most individuals from Cumberland Sound and Eastern Hudson Bay and Sanikiluaq, and all individuals from Bear Island in northern James Bay (Figure S1). Through a PCA using exclusively the 84 individuals from this third cluster (Figure 2b) and K=5 and K=6 admixture analyses, we identified three additional populations. The Cumberland Sound population consisted of individuals sampled from Cumberland Sound and one individual from Iqaluit. The Hudson Bay-Strait Complex population comprised all individuals from Western Hudson Bay, Ungava Bay and Bear Island, nine of the ten individuals sampled near Iqaluit (Baffin Island), most beluga harvested in Eastern Hudson Bay and the Belcher Islands, and the one beluga from the Strait of Belle Isle (Figure 2b). The Little and Great Whale Rivers population (named following Montana et al. 2024), consisted of individuals sampled in Eastern Hudson Bay, specifically from the Little Whale River and around Kuujjuarapik at the mouth of the Great Whale River. It additionally included three individuals from Eastern Hudson Bay for which we had no specific sampling location. Ancestral admixture at K=7 did not result in further population distinction and instead showed admixture within the Hudson Bay-Strait Complex population (Figure S2). While fine structure has been documented in the Hudson Bay-Strait Complex in other studies (Parent et al., 2023), our dataset in the present study does not distinguish subgroups within this population, which could be due to sample size. Since demographic history and effective population size analysis rely on distinctive populations, we treated this complex as one genetic population.

**Figure 2.**
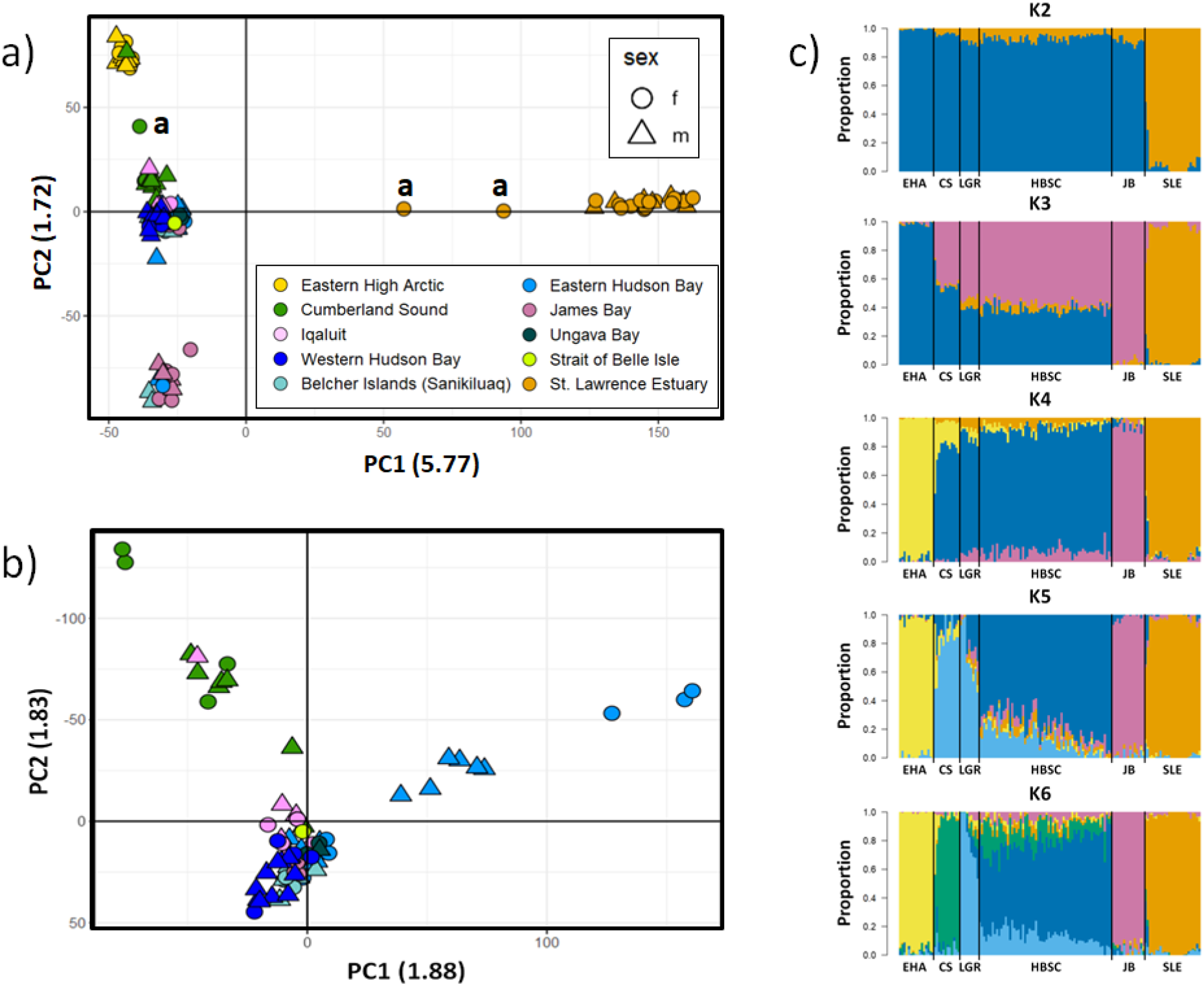
a) PCA showing the population structure of all sampled individuals (n=140). Individuals treated as “admixed” for further analysis are labeled in the PCA with an “a”. b) PCA of population structure within the largest, highly heterogeneous cluster from Figure 2a (n=84). c) Admixture plot with K=2–6. EHA=Eastern High Arctic, CS=Cumberland Sound, LGR=Little and Great Whale Rivers, HBSC=Hudson Bay-Strait Complex, JB=James Bay and SLE=St. Lawrence Estuary.

The St. Lawrence Estuary population was the most distinct with *F*_ST_ values > 0.08 compared to all other populations (highest *F*_ST_ of 0.12 was between St. Lawrence Estuary and the Eastern High Arctic) (Table 1). The St. Lawrence Estuary population was also the most inbred population with a mean ROH over 550 Mb; all other populations had a mean ROH ∼ 200 Mb (Figure 3). The Eastern High Arctic and James Bay populations had intermediate *F*_ST_ when compared with the other Arctic populations. The Cumberland Sound, and Little and Great Whale Rivers populations, as well as belugas from the wider Hudson Bay-Strait Complex had the lowest *F*_ST_ values, showing less divergence compared to the other populations (*F*_ST_ range: 0.0083 - 0.0109; Table 1).

**Figure 3.**
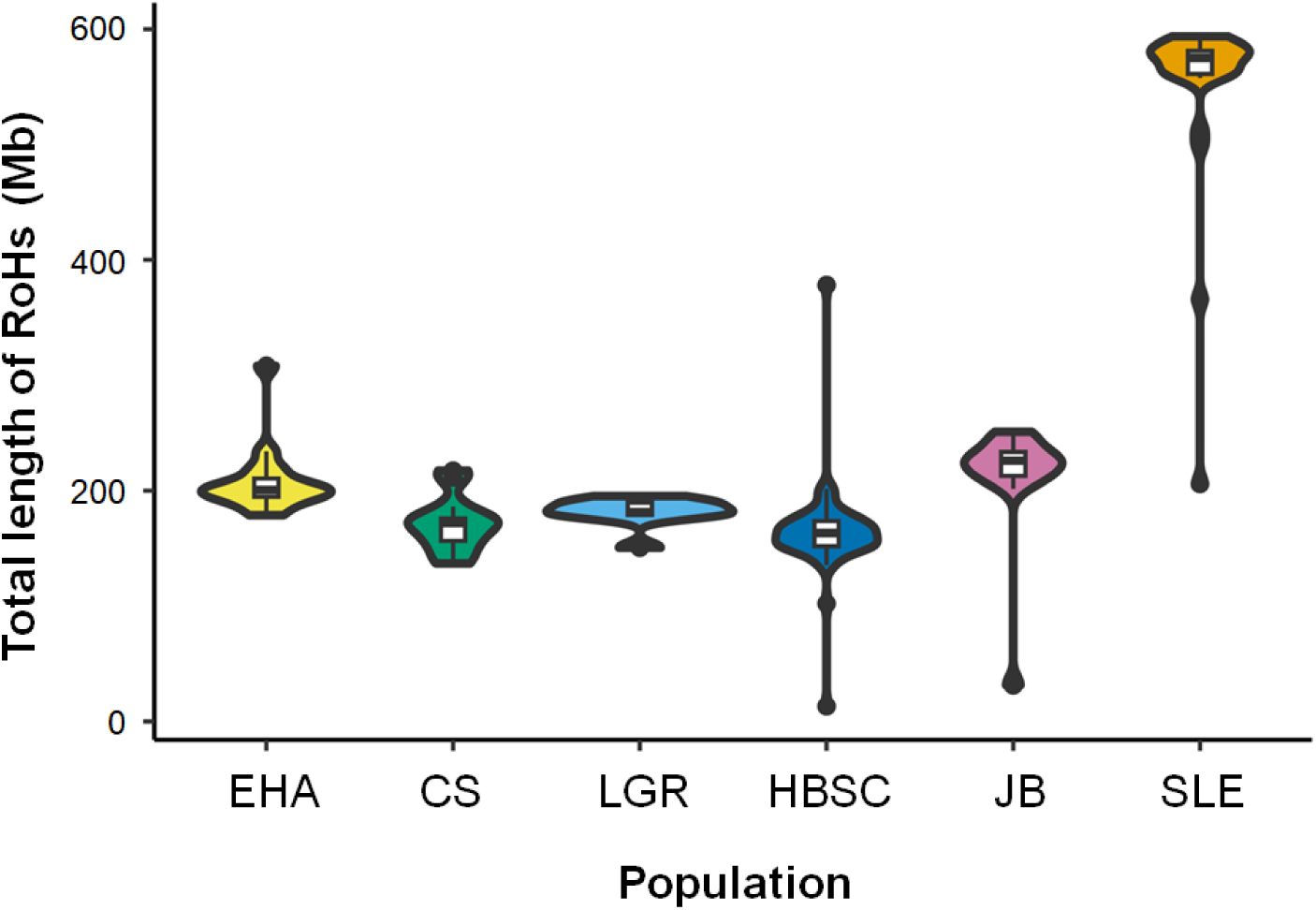
Runs of homozygosity estimates for each population (EHA=Eastern High Arctic, CS=Cumberland Sound, LGR=Little and Great Whale Rivers, HBSC=Hudson Bay-Strait Complex, JB=James Bay, SLE=St. Lawrence Estuary).

**Table 1.** Weighted pairwise *F*_ST_ between genetically distinct populations. EHA=Eastern High Arctic, CS=Cumberland Sound, HBSC=Hudson Bay-Strait Complex, LGR=Little and Great Whale Rivers, JB=James Bay and SLE=St. Lawrence Estuary.

| Pairwise $F_{ST}$ | EHA | CS | HBSC | LGR | JB |
| --- | --- | --- | --- | --- | --- |
| **CS** | 0.0206 |  |  |  |  |
| **HBSC** | 0.0238 | 0.0083 |  |  |  |
| **LGR** | 0.0362 | 0.0203 | 0.0109 |  |  |
| **JB** | 0.0497 | 0.0359 | 0.0269 | 0.0426 |  |
| **SLE** | 0.1217 | 0.1055 | 0.0876 | 0.1129 | 0.1191 |

### Effective Population Size & Demographic History

Our *SMC++* results showed synchronous trajectories in *N*_e_ for all populations until the middle of the last glacial period, some 50,000 years ago (Figure 4, see Figure S3 for all iterations). Afterwards, we observed a decline in *N*_e_ for the St. Lawrence Estuary, the Eastern High Arctic, the James Bay and the Little and Great Whale Rivers populations. In contrast, the *N*_e_ for the Cumberland Sound and Hudson Bay-Strait Complex populations remained largely stable, with the Hudson Bay-Strait Complex population showing a slight increase towards the end of the glacial period. While *N*_e_ for the Eastern High Arctic, James Bay, and Little and Great Whale Rivers populations continued to decline, the St. Lawrence Estuary population experienced a sharp decrease at first, followed by a gradual increase in effective population size, starting around 10,000 years ago.

**Figure 4.**
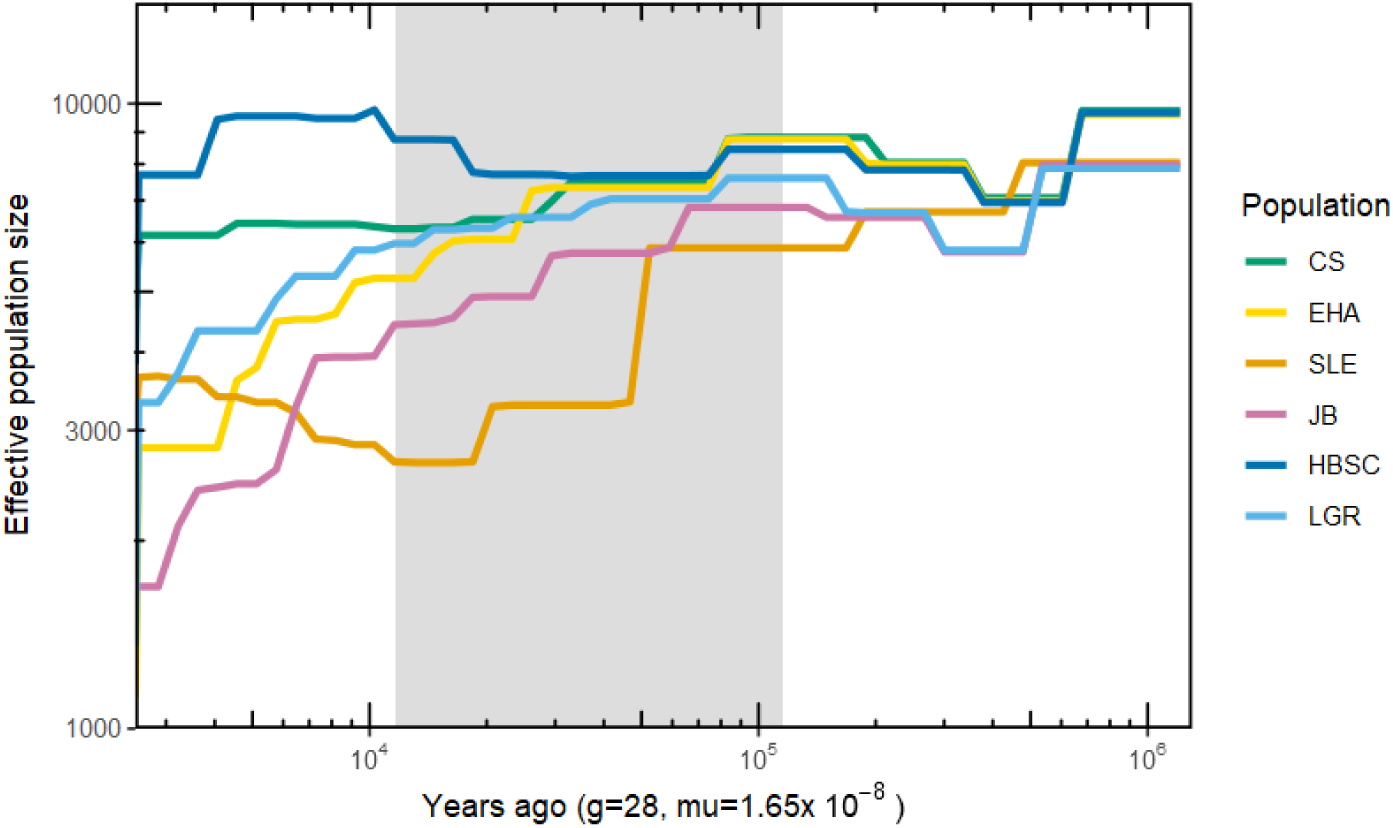
Changes in beluga whale effective population size (*N_e_*) over time shown by the median of 20 SMC++ iterations per population. The grey bar marks the last glacial period (115 – 11.7 thousand years ago) and axes are log10-scaled. CS=Cumberland Sound, EHA=Eastern High Arctic, SLE=St. Lawrence Estuary, JB=James Bay, HBSC=Hudson Bay-Strait Complex, LGR=Little and Great Whale Rivers.

Recent (3,000 years ago to present) demographic history reconstructions using *GONE* differed between the six populations (Figure 5; Figure S4). We observed an increase in *N*_e_ of three populations in the last 500 -- 1,000 years. The Eastern High Arctic population showed a decline in *N*_e_ around 2,000 years ago, followed by an increase over the past 1,000 years (Figure 5). The St. Lawrence Estuary population maintained a stable *N*_e_ over most of the period but showed an increase in *N*_e_ starting 500 years ago, with a stabilization and possible decrease in *N*_e_ over the past 100 years. The Hudson Bay-Strait Complex population showed two substantial increases in N_e_, the first one 2,500 years ago and the second one 1,000 years ago. The *N*_e_ for the James Bay population decreased until 1,500 years ago, to increase slightly up until 500 years ago, at which point the model estimated another drop in *N*_e_ numbers. The *N*_e_ for the Cumberland Sound and Little and Great Whale Rivers populations decreased over the last 1,000 years. In the case of the Cumberland Sound population, the decrease was gradual and followed a period of increased *N*_e_, while for the Little and Great Whale Rivers population, the decrease in *N*_e_ was more gradual over the last 2,500 years until 750 years ago where we noted a steep drop in *N*_e_, which aligned with the medieval warm period.

**Figure 5.**
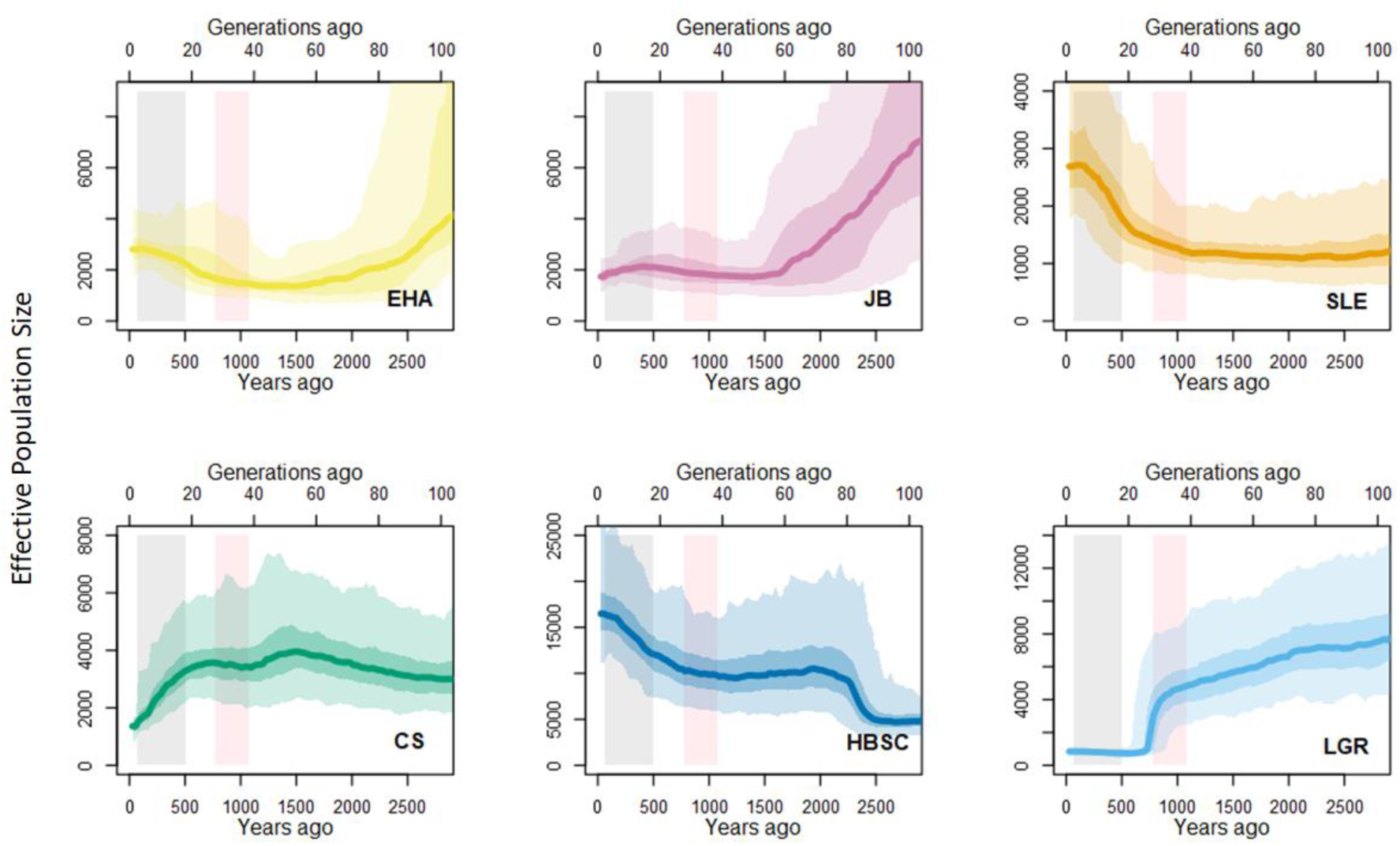
Demographic history reconstruction of the past 100 generations using a linkage disequilibrium-based method *GONE*. Results are presented as the median (solid line) and 95% confidence interval (light shading). Darker shading shows the interquartile range. EHA=Eastern High Arctic, JB=James Bay, SLE=St. Lawrence Estuary, CS=Cumberland Sound, HBSC=Hudson Bay-Strait Complex, and LGR=Little and Great Whale Rivers. The pink shaded area represents the Medieval Warm Period (from 950 to 1250 years ago). The grey shaded area represents the time of commercial hunting from early 16^th^ century (500 years ago) to the end of commercial whaling in Canada (60 years ago).

The estimates for the contemporary *N*_e_ suggest the Eastern High Arctic and Hudson Bay-Strait Complex populations have the highest contemporary *N*_e_ estimates, at 2,193 and 3,108 respectively (Table 2 & S6). The remaining northern populations all have *N*_e_ estimates below 400, with the lowest estimate in the Little and Great Whale Rivers population at *N*_e_ = 123. The *N*_e_ of the St. Lawrence Estuary population varied depending on whether admixed individuals were included or not, and the degree of relatedness between individuals (Table S6). Estimating *N*_e_ using all samples from St. Lawrence Estuary resulted in an *N_e_* of 989 or 1,891 when conducted without admixed individuals. Retaining only 10 samples of the least related individuals (see methods) resulted in a *N*_e_ estimate of 185 or 1,995 if conducted without admixed individuals.

**Table 2.** Effective population size estimates of the six genetic populations. “*n*” represents the number of genomic samples. “*N*” represents the estimated size of the entire population based on aerial surveys, which are extracted from the most recent COSEWIC or DFO status reports (COSEWIC, 2020; Sauvé et al., 2024; Tinker et al., 2024). The HBSC abundance estimate *N* represents the combined estimates for Western Hudson Bay beluga and the Eastern Hudson Bay - Belcher Islands stock and thus, include beluga from LGR. “*N*_e_” is the average estimated effective population size, based on 10 iterations of a random subsampling of 10,000 SNPs from our data. The average lower and upper 95% confidence limits (CI) are for *N*_e_ estimates of each population. “low kinship only” refers to the estimates where we only used ten individuals with the lowest degree of relatedness for estimation of Ne.

| Population | $n$ | $N$ | $N_e$ | Lower 95% CI | Upper 95% CI |
| --- | --- | --- | --- | --- | --- |
| EHA | 16 | 21,213 | 2,193 | 1,773 | 2,896 |
| CS | 12 | 1,090 | 346 | 333 | 361 |
| HBSC | 62 | 58,292 | 3,108 | 2,922 | 3,319 |
| LGR | 9 | Na | 123 | 120 | 127 |
| JB | 15 | 11,455 | 266 | 259 | 274 |
| SLE | 26 | 1,850 | 989 | 937 | 1,047 |
| SLE (low kinship only) | 10 | 1,850 | 185 | 179 | 192 |

## Discussion

We assessed the demographic history of the beluga whale in a rapidly warming environment in eastern Canada. Recent trends and contemporary effective population size estimates suggest that some beluga whale populations may be more vulnerable to losses of genetic diversity than their census size might indicate. Many of the identified populations have lower effective population sizes than they had 2,500 years ago, with two of the smallest three populations (Cumberland Sound and Little and Great Whale Rivers) showing a continued decline starting 500 to 1,000 years ago. The St. Lawrence Estuary population, the third small population, showed an increase in *N*_e_ until about 100 years ago. The effective population sizes of the larger populations (Eastern High Arctic and Hudson Bay-Strait Complex) have also increased in recent history. Contemporary *N*_e_ estimates were small, in the hundreds, for all populations except for the two populations with the largest population sizes based on aerial surveys, which had *N*_e_ values above 2,000.

### Population dynamics and deep time demographic history

We documented six genetically distinct populations, consistent with the findings of Montana et al. (2024) who had access to a more extensive and spatially resolved dataset. Given the strong consensus on population structure, we based our demographic analyses on these well-defined genetic populations.

Our reconstruction of long-term demographic change suggests that the ancestors of current populations had a shared demographic history until approximately 50,000 years ago, near the middle of the last glacial period. This indicates that the current beluga populations began to diverge during the last glacial period, which is consistent with the hypothesis that the St. Lawrence Estuary population was separated from the Northern Beluga population around that time (Harington, 1977). The onset of demographic divergence between the populations also marks the beginning of a decrease in effective population size for all populations except Cumberland Sound and Hudson Bay-Strait Complex (Figure 4). Ice masses reached their last maximum global extent approximately 18,000 years ago (Martinson et al., 1987). Our results suggest that our genetic populations were isolated from each other while ice masses were growing, reducing the effective population sizes in all but the Hudson Bay-Strait Complex population. The decline in *N*_e_ likely resulted from the split of one large population into multiple smaller ones. The *N*_e_ of the Eastern High Arctic, the James Bay and the Little and Great Whale Rivers populations continued to decline gradually after the last glacial maximum. The sharp decrease in *N*_e_ that was observed for St. Lawrence Estuary beluga between 50,000 and 10,000 years ago, which was followed by an increase after 10,000 years ago, might be attributable to this population migrating to the Gulf of St. Lawrence as this area opened up some 15,000 years ago (Occhietti et al., 2001). This area increased in size with the appearance of the Champlain Sea, an inland sea connected to the Atlantic Ocean covering most of the St. Lawrence lowlands between 13,000 and 10,000 years ago. Fossils of beluga whales have been discovered in the Champlain Sea (Harington, 1977). The extent of the beluga habitat grew between 15,000 and 10,000 years ago, and was much larger than it is today, potentially offering this population new favorable habitat which allowed it to expand in numbers during the warmer post glacial period.

### Recent demographic history patterns

One significant threat to belugas in recent centuries has come from human hunting associated with commercial whaling in the post-colonial era, starting in the early 16^th^ century. While early commercial whaling did not often target belugas and was primarily focused on bowhead and right whales (Loewen, 2009), belugas were intensively harvested in some regions. Commercial beluga whale harvest in Cumberland Sound, the Little Whale and Great Whale rivers in the Eastern Hudson Bay, and in the St. Lawrence Estuary resulted in intense exploitation of these populations (Béland, 1996; Mitchell and Reeves, 1981). We therefore might expect to see substantial declines in the effective population sizes of these populations, coinciding with commercial whaling activities. However, our reconstructions of recent effective population size changes suggest that two of the most threatened stocks, the Little and Great Whale Rivers and the Cumberland Sound populations, initiated their decline in effective population sizes before the beginning of the commercial whaling era. The original decline in *N*_e_ for Cumberland Sound, which began approximately 1,500 years ago, appeared to have stopped during the medieval warm period, a time of local high temperatures in Earth’s Northern Hemisphere (Hughes and Diaz, 1994; Zhou et al., 2011). While little is known about the impact of this warming period on the climate in the Canadian Arctic, our results suggest that this period may have affected effective population size in at least three of our six populations. While this event coincided with an increase in the effective population size in the Hudson Bay-Strait Complex and a halt in the decline of the Cumberland Sound Population, the medieval warm period was also correlated with the most significant and most extreme drop in the effective population size in the Little and Great Whale Rivers population. It is possible that retreating sea ice during the medieval warm period has resulted in this population getting exposed more to predators such as killer whales; however, if this is the explanation for this rapid decline, we would have expected there to be similar occurrences in other populations nearby, such as the Hudson Bay-Strait Complex or the James Bay populations. This was not the case.

The Little and Great Whale Rivers population is the only population that experienced decline that may be attributable to commercial harvest, although a more slowly progressing decline was also observed in the Cumberland Sound population. This decline is most pronunced during the commercial whaling period; however, it also begins earlier, at the same time as the decline in the Little and Great Whale Rivers population and might therefore be causally linked. It is still possible that the decline in the Little and Great Whale River population is linked to commercial whaling. Recent negative effects might be difficult to detect against the backdrop of recent decline.

The only northern population that does not appear to have been affected by either the environmental changes during the medieval warm period or the commercial whaling of the last centuries is the James Bay population, which has remained largely stable over the last 1,500 years. Only a slight and gradual decline in *N*_e_ was detected since commercial whaling began. The fact this population has not been subject to commercial whaling and remains in James Bay year-round (Bailleul et al., 2012) has likely limited both gene flow into this population and harvest removals. Nevertheless, it is possible that at least some James Bay beluga might be taken out by whaling in Hudson Bay. While there is no evidence of commercial hunting in James Bay itself, many samples from this genetic population have been collected outside James Bay, at Kuujjuarapik and Belcher Island (figure S1, table S1). It is therefore conceivable that hunting along the Eastern Hudson Bay could also affect the James Bay population. This could explain the slight dip we detect at the onset of the commercial whaling period in their demographic history.

The suggested increase in the effective population size of the third most threatened population, the St. Lawrence Estuary, over the past centuries was unexpected given that commercial harvest removed thousands of individuals during this period (Béland, 1996). Gene flow from Arctic populations into the St. Lawrence Estuary during this period could help account for the observed pattern. In genetically segregated populations, sudden occurrence of gene flow can rapidly increase effective population size without affecting total abundance (Novo et al., 2023). There is evidence for at least episodic influxes of Arctic beluga into the St. Lawrence Estuary from the literature (Vladykov, 1944). The relative proximity of wintering areas of some of the Arctic populations in southern Labrador (Bailleul et al., 2012), reports of vagrant beluga each summer in Newfoundland waters, the northwestern and southern Gulf of St. Lawrence, Scotian Shelf or eastern U.S. (Curren and Lien, 1998; R. Michaud, unpublished data), with some identified genetically or from their contaminant loads as coming from the Arctic, provide additional evidence for potential gene flow coming from the northern populations (Lesage et al., 2024). The genetic identity of the Strait of Belle Isle beluga in this study (attributed to the Hudson Bay-Strait Complex population), although beach-cast and therefore with a degree of uncertainty about its location while still alive, further support the presence of Arctic beluga in southern waters. The two St. Lawrence Estuary individuals that were admixed with the Hudson Bay-Strait Complex population further support exchanges with Arctic populations. It is important to note that low levels of gene flow like this may not significantly mitigate inbreeding stress in this population. Studies on inbreeding and mutation load have supported the conclusion that this population is inbred (DFO Canadian Science Advisory Secretariat Science Advisory Report, 2024; Orton et al., 2025), even if there might be more gene flow than previously expected. Over the past 100 years, *N*_e_ appears to have stabilized and possibly decreased for St. Lawrence Estuary beluga, suggesting a reduced influx of Arctic animals. The population decline observed for St. Lawrence beluga and some of the eastern Arctic populations (e.g., Cumberland Sound, Eastern Hudson Bay), combined with the warming climate, might have contributed to limit gene flow with this southern population over the past century.

While an increase in *N*_e_ was noted both for the Eastern High Arctic and Hudson Bay-Strait Complex populations during the last 1000 years, reasons behind these increases are likely to be different for the two populations. Left largely undisturbed, the Eastern High Arctic population seemed to have slowly recovered from the decline in *N*_e_ following the last glacial period, probably furthered by suitable environmental conditions and lack of natural or anthropogenic disturbances. Other factors might explain the rapid increase in the Hudson Bay-Strait Complex population. Studies with larger sample sizes and higher resolution have detected finer structure in this population (Parent et al., 2023), and our admixture analysis reveals that the Hudson Bay and Strait Complex population contains a diverse range of genetic combinations from different ancestries (Figure 2c). The two increases we see in our demographic history reconstruction over the last centuries could indicate instances of mergers between previously more distinct populations. This could result in a large genetically distinct cluster containing substructure from previously distinct populations with large spikes in their demographic history which indicate the onset of gene flow between them.

### Contemporary N_e_

Both historical and contemporary *N*_e_ can help inform conservation management. Modern conservation and management efforts have started to consider *N*_e_ as a measure for the adaptive capacity of wildlife populations. In 2022 the Convention on Biological Diversity formulated the Kunming-Montreal Global Biodiversity framework, which established an *N_e_* of 500 as a baseline for the measure of the genetic health of a population (Franklin, 1980; Jamieson and Allendorf, 2012). A population with a *N*_e_ of at least 500 is often considered healthy, with a capacity to maintain adaptive potential (Hoban et al., 2020; Jamieson and Allendorf, 2012). However, this metric should be considered in light of the population’s demographic history, including periods of bottlenecks and expansions. The large contemporary *N*_e_ estimated for the Eastern High Arctic and Hudson Bay-Strait Complex populations and the continued increase in *N*_e_ suggest these populations are stable in their genetic diversity. This could be indicative of a steady or large adaptive capacity, which may help provide resilience to future changes such as a warming climate. In case of the Eastern High Arctic, this will likely be a requirement for this population to persist in the future, given that this region is predicted to require a higher degree of genetic adaptation to environmental changes over the next decades, given current climate change models (De Greef et al., 2025). The lower values of *N*_e_ for the other Arctic populations suggest these populations may be at greater risk of genetic diversity loss due to genetic drift, potentially leading to lower adaptive capacity. Most notably, the Little and Great Whale Rivers population had a substantial decline in *N*_e_ in its recent past, as well as the lowest contemporary *N*_e_. The observed low *N*_e_ has to be considered carefully given the low sample size we have for this population. However combined with the recent decline in this population it suggests that this population is most susceptible to further declines and future environmental changes.

The contemporary *N*_e_ of the St. Lawrence Estuary population varied from 185 to 1,995 individuals depending on underlying assumptions about the samples used (Table S6). These estimates are in line with previous estimates in this population (e.g., Best, 2022). The St. Lawrence Estuary population had longer ROHs compared to the other beluga populations, indicating elevated levels of inbreeding and high risk of genetic diversity loss through genetic drift.

The population with the next highest ROH value is the James Bay population, although it is not significantly higher than the remaining populations. This population has an estimated abundance exceeding 10,000 beluga (Sauvé et al. 2024), indicating little reason for concern. However, its contemporary *N*_e_ is below 500 and similar to the *N*_e_ of the Cumberland Sound population. The demographic history reconstruction of James Bay beluga reveals no significant changes in *N*_e_ in the recent past, which suggests that the genetic diversity of this population has remained relatively stable over the last few centuries and may not have been heavily affected by climate and anthropogenic factors. However, recent projections for Arctic beluga populations puts the James Bay population at higher need for adaptation to a changing environment than most other beluga populations (De Greef et al., 2025). Despite its demographic stability over the last centuries, its low *N*_e_ in an environment predicted to change strongly in the near future indicates a population more at risk than its pure census data would indicate.

### Implications

The conservation of genetically distinct populations is important for many reasons. Among the most important is the long-term survival and adaptability of species in the face of environmental change. Each genetic population may harbour genetically distinctive allele frequencies, which are not just a record of the past but a resource for the future, providing the raw material upon which natural selection can act as environmental conditions shift. Our findings in beluga whale genomics reveal a complex demographic history shaped by both natural climatic fluctuations, such as post-glacial habitat changes, and human influences, including commercial hunting. These forces have not acted uniformly across the range of Canada’s belugas. As a result, some populations now exhibit decreased levels of genetic diversity and small effective population sizes, which in turn reduce their capacity to adapt to ongoing and future environmental pressures. Other populations have shown high genetic diversity and signs of demographic recovery, suggesting a difference in potential for resilience among groups. Maintaining genetic diversity within, and gene flow between these populations is essential for enhancing species-wide resilience. The disparities in genetic diversity, recent demographic trajectories, and effective population size among beluga populations in eastern Canada emphasizes the inadequacy of one-size-fits-all conservation measures. Each population’s distinct evolutionary history and current status requires specific management strategies that address local conditions. Conservation planning should therefore integrate genetic data in addition to ecological considerations to facilitate the success of these animals in the future.

## Acknowledgements

We are grateful to Nunavik and Nunavut Inuit hunters and stakeholders, as well as the many partners involved in the long-term carcass recovery program in the St. Lawrence Estuary for providing the samples used in this study. Additional thanks goes to Dr. Matt Thorstensen for his help with the *SMC++* analysis. This work was supported by the Natural Sciences and Engineering Research Council (NSERC), Fisheries and Oceans Canada (DFO), the University of Manitoba and the Digital Research Alliance of Canada.

## Notes

### Competing Interest Statement

The authors have declared no competing interest.

