## Supplemental Material for "Exploring contemporary and historic effective population sizes in Atlantic beluga whale populations"

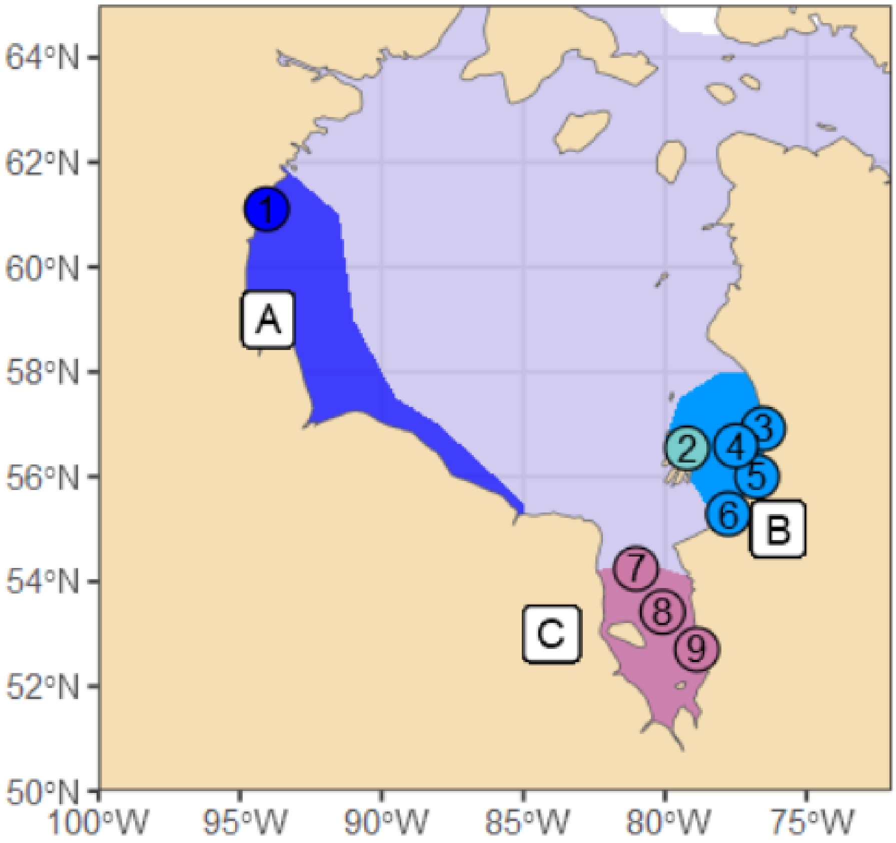


**Figure S1: Map of Hudson Bay & James Bay sampling sites.** The colors represent the different stocks as described by COSEWIC (2020): Western Hudson Bay (A, blue), Eastern Hudson Bay (B, light blue) & James Bay (C, purple). Teal shows the Sanikiluaq sampling site. The numbers represent the sampling sites: (1) Arviat, (2) Sanikiluaq, (3) Nastapoka River, (4) NA (no detailed sampling information, therefore allocated to the center of the Eastern Hudson Bay summer habitat), (5) Little Whale River, (6) Kuujjuarapik, (7) Bear Island, (8) NA (no detailed sampling information, therefore allocated to the center of the James Bay summer habitat) and (9) Pointe de Repentigny.

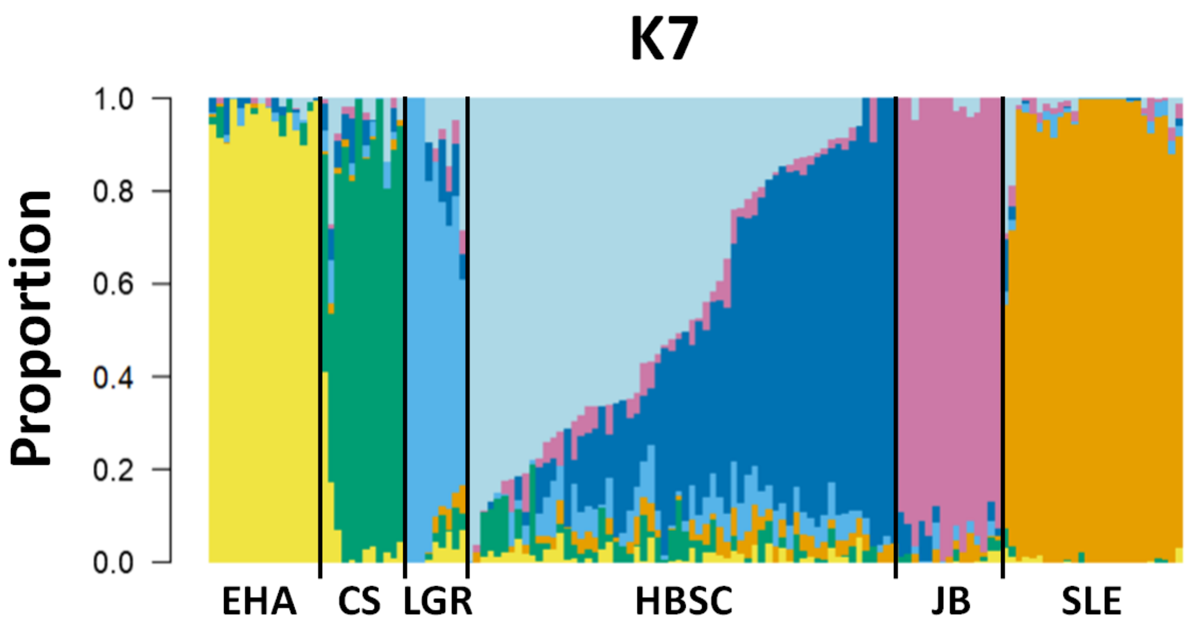


**Figure S2: Admixture plot for K=7.** EHA=Eastern High Arctic, CS=Cumberland Sound, LGR= Little and Great Whale Rivers, HBSC=Hudson Bay-Strait Complex, JB=James Bay, SLE=St. Lawrence Estuary. K=7, compared to K=6, creates a genetic continuum within HBSC, instead of a new distinct population. The order of this progression does not follow any discernable geographic pattern.

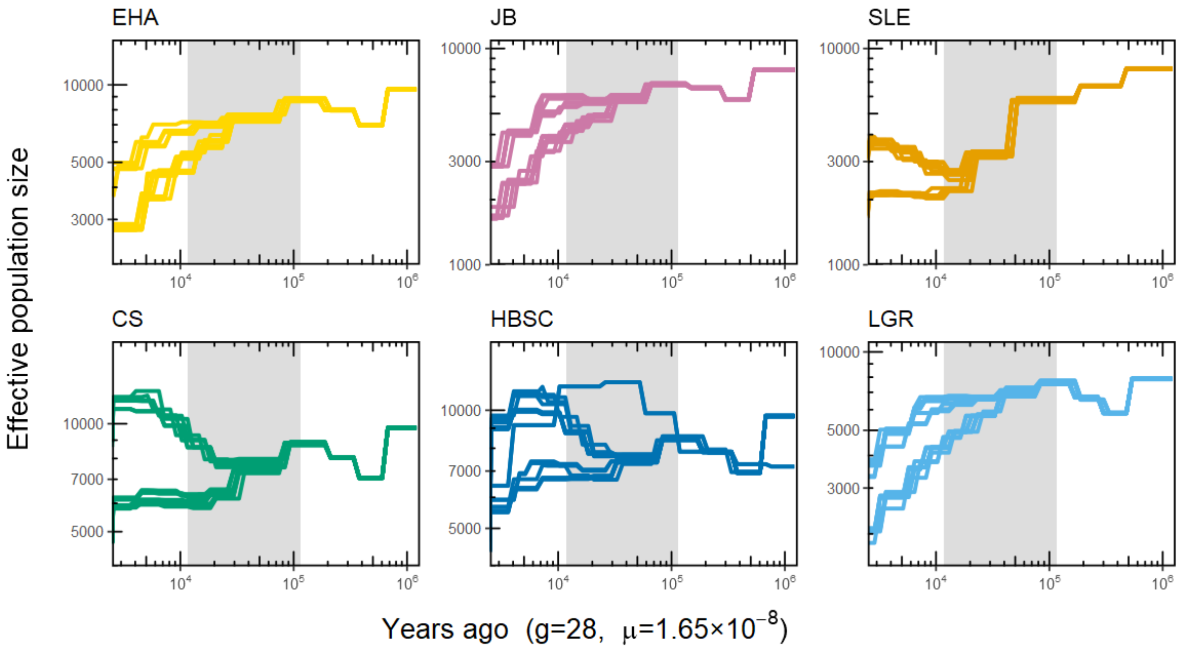


**Figure S3: Plots of all 20 SMC++ iterations per population.** EHA=Eastern High Arctic, JB=James Bay, SLE=St. Lawrence Estuary, CS=Cumberland Sound, HBSC=Hudson Bay and Strait Complex, LGR=Little and Great Whale River. The grey bar marks the last glacial period.

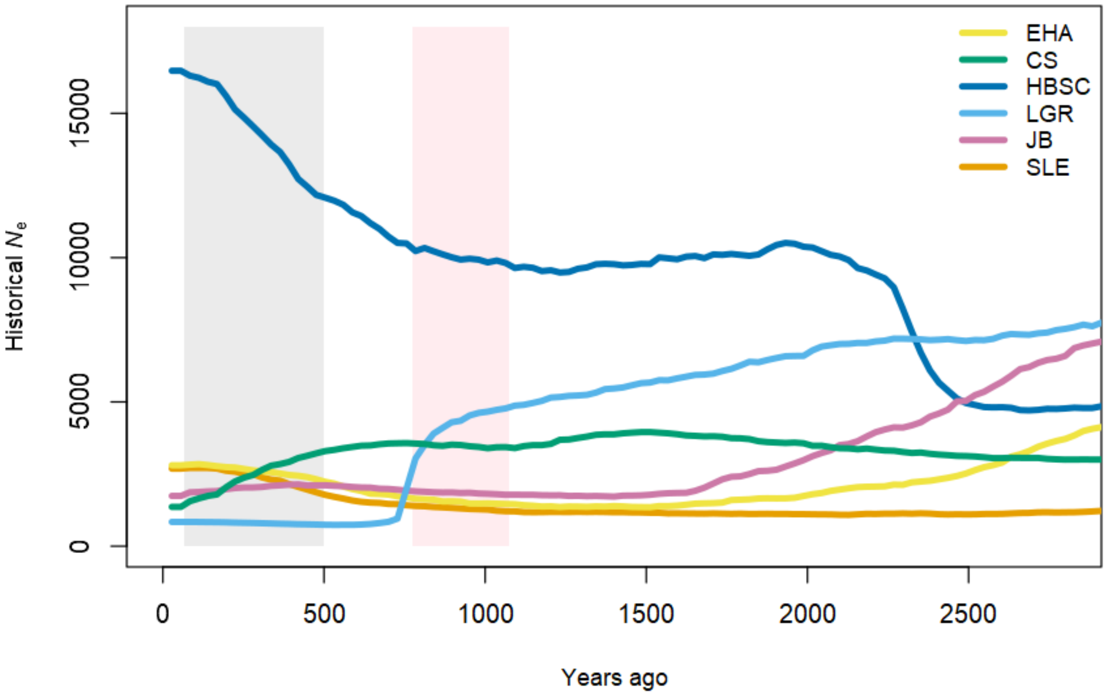


**Figure S4: Demographic history reconstruction of the last 100 generations.** Eastern High Arctic (yellow), Cumberland Sound (green), Hudson Bay-Strait Complex (blue), Little and Great Whale Rivers (light blue), James Bay (purple) and St. Lawrence Estuary (orange). The pink shaded area represents the Medieval Warm Period (from 950 to 1250 years ago). The grey shaded area represents the time of high commercial hunting from early 16th century (500 years ago) to the end of commercial whaling in Canada (60 years ago). We used 28 years/generation as generation time.

**Table S1: Distinct genetic populations for SMC++.** The sites column lists all the sample sites represented within this population. If a site is listed in more than one population, it means that we had multiple individuals from that site and not all individuals corresponded to the same population during the population structure analysis. NA in this column means that the exact sampling site was not known. These individuals are from the shores somewhere along the Eastern Hudson Bay, James Bay or in the St. Lawrence estuary. The number in brackets shows how many individuals from each site correspond to any given population. A lower case "a" inside the bracket indicates that this individual is admixed and does not cluster fully with that population. The number of individuals column shows how many individuals we have for each population. The Distinguished lineage column shows which individual was chosen to represent the distinguished lineage for the SMC++ analysis.

| Population | Sites | Number of Individuals | Distinguished lineage |
|---|---|---|---|
| **Eastern High Arctic** | Cunningham Inlet (15)<br>Pangnirtung (1) | 16 | EHA_Cunningham_Inlet_104 |
| **Cumberland Sound** | Pangnirtung (10)<br>Pangnirtung (1, a)<br>Iqaluit (1) | 12 | CS_Pangnirtung_1022 |
| **Hudson Bay- Strait Complex** | Iqaluit (9)<br>Kuujjuarapik (4)<br>Nastapoka River (4)<br>Umiujaq (1)<br>Sanikiluaq (13)<br>Bear Island (3)<br>Pointe De Repentigny (1)<br>Strait of Belle Isle (1)<br>Cap Qualirusiq (1)<br>Leaf Bay (2)<br>Arviat (15)<br>NA (7)<br>Pangnirtung (1) | 62 | HB_Sanikiluaq_30 |
| **Little and Great Whale Rivers** | Kuujjuarapik (4)<br>Little Whale River (2)<br>NA (3) | 9 | EHB_Kuujjuarapik_3812 |
| **James Bay** | Kuujjuarapik (3)<br>Sanikiluaq (2)<br>Pointe De Repentigny (9)<br>NA (1) | 15 | JB_Pointe_De_Repentigny_975 |
| **St. Lawrence Estuary** | Baie-Sainte-Catherine (1)<br>Baie-des-Sables (1)<br>Cacouna (1)<br>Cap-aux-Oies (1)<br>Grand-Métis (1)<br>Île aux Basques (1)<br>Isle Verte (1)<br>Matane (2)<br>Rimouski (1)<br>Rimouski Est (1, a)<br>Grosses-Roches (1, a)<br>Rivière-du-Loup (1)<br>Saint-Ulric (3)<br>Sainte-Flavie (1)<br>NA (9) | 26 | SL_Matane_4293 |

**Table S2: Summary of base filter steps for the genomic data used for all analyses.** The filter steps are listed as well as the command and parameter used. SNP column denotes how many SNPs are left after each step. The n column shows the number of individuals.

| Filter | Parameter | SNPs left | n |
|---|---|---|---|
| **None** | - | 17,203,219 | 146 |
| **Remove indels** | --remove-indels | 13,069,604 | 146 |
| **Only keep sites flagged with "PASS"** | --remove-filtered-all | 7,224,915 | 146 |
| **Only keep sites with quality score >50** | --minQ 50 | 7,013,152 | 146 |
| **Remove sites with missingness >25%** | --max-missing 0.75 | 7,008,512 | 146 |
| **Remove non-biallelic sites** | --max-alleles 2<br>--min-alleles 2 | 6,995,734 | 146 |
| **Remove scaffolds <100kbp & sex chromosomes** | --not-chr… | 6,800,997 | 146 |
| **Remove close kin individuals (pi-hat>0.4) and individuals with missingness>30%** | --remove-indv… | 6,800,997 | 140 |

**Table S3: Summary of filter steps for the genomic data used for population structure analyses.** The filter steps are listed as well as the command and parameter used. SNP column denotes how many SNPs are left after each step. The n column shows the number of individuals.

| Filter | Parameter | SNPs left | n |
|---|---|---|---|
| **After base filter** | - | 6,800,997 | 140 |
| **Remove sites out of Hardy-Weinberg equilibrium** | --hardy<br>--exclude-positions | 6,797,823 | 140 |
| **Remove alleles with lower frequency than 5%** | --maf 0.05<br>--max-maf 0.95 | 3,091,381 | 140 |
| **Prune variants within 1000bp of each other** | --thin 1000 | 1,081,765 | 140 |

**Table S4: Summary of filter steps for the genomic data used for effective population size and demographic history analysis.** The filter steps are listed as well as the command and parameter used. The "Number of Samples" line shows how many SNPs are left after each step and the number of individuals in brackets.

| Filter | Parameter | SNPs (samples) |
|---|---|---|
| **After base filter** | - | 6,800,997 (n=140) |
| **Only keep chromosome aligned sites** | --not-chr… | 6,797,654 (n=140) |
| **Remove sites out of Hardy-Weinberg equilibrium** | --exclude-positions | 6,794,859 (n=140) |

**Table S5: Parameter settings for GONE analyses.**

| Parameter | Setting | Description |
|---|---|---|
| PHASE | 2 | Used for unphased data (0=pseudohaploids, 1=phased, 0=unphased) |
| cMMb | 1 | Average rate of recombination assumed in CentiMorgans per megabase |
| DIST | 1 | Type of genetic distance correction (0=no correction, 1=Haldane correction, 2=Kosambi correction) |
| NGEN | 2000 | Number of generations for which linkage data is obtained in bins |
| NBIN | 400 | Number of bins. |
| MAF | 0.0 | Minor allele frequency cut off, 0 is recommended. |
| ZERO | 1 | Determines how to deal with SNPs with zero value (0=remove, 1=allow) |
| maxNCHROM | -99 | Maximum number of chromosomes used (-99 = all chromosomes are used) |
| maxNSNP | 20,000 | Maximum numbers of SNPs/chromosome used |
| hc | 0.05 | Maximum value of recombination rate being analysed. |
| REPS | 500 | Number of replicates to run per analysis |
| threads | -99 | Number of threads used (-99 uses all possible processors) |

**Table S6.** Population size estimates of our populations. “*n*” represents the number of genomic samples. The columns "SNPs” shows how many SNPs were kept after all the filtering steps. “*N*” represents the estimated census size of the entire population, which are extracted from the most recent COSEWIC or DFO status report (COSEWIC 2020; Sauvé u. a. 2024; Tinker u. a. 2024). “$N_e$” is the average estimated effective population size, based on 10 iterations of a random subsampling of 10,000 SNPs from our data. Upper and Lower 95%-CI shows the average upper and lower 95% confidence interval for $N_e$ for each population. "w/o admixed" indicate the estimates that were done while excluding admixed individuals. "low kin" refers to the estimates where we only used the ten individuals with the lowest degree of relatedness.

| **Pop** | ***n*** | **SNPs** | ***N*** | **$N_e$** | **Lower 95% CI** | **Upper 95% CI** |
|---|---|---|---|---|---|---|
| **EHA** | 16 | 2,156,788 | 21,213 | 2,193 | 1,773 | 2,896 |
| **CS** | 12 | 2,239,053 | 1,090 | 346 | 333 | 361 |
| **CS w/o admixed** | 11 | 2,132,238 | 1,090 | 288 | 277 | 301 |
| **HBSC** | 62 | 789,167 | 58,292 | 3,108 | 2,922 | 3,319 |
| **LGR** | 9 | 1,598,774 | Na | 123 | 120 | 127 |
| **JB** | 15 | 925,442 | 11,455 | 266 | 259 | 274 |
| **SLE** | 26 | 658,098 | 1,850 | 989 | 937 | 1,047 |
| **SLE w/o admixed** | 24 | 791,979 | 1,850 | 1,891 | 1696 | 2,138 |
| **SLE (low kin)** | 10 | 1,328,142 | 1,850 | 185 | 179 | 192 |
| **SLE (low kin) w/o admixed** | 10 | 1,502,548 | 1,850 | 1,995 | 1414 | 4,310 |

**Expanded discussion on the effective population size of the St. Lawrence Estuary population:**

$N_e$ estimates are greatly influenced by different model inputs regarding genetic outliers (admixed individuals) and the degree of relatedness between the individuals. In the Cumberland Sound population, including admixed individual results in an increase of $N_e$ while in St. Lawrence Estuary population, excluding admixed individuals resulted in a very strong increase in $N_e$ estimates. The St. Lawrence Estuary estimates stand out as unusual because their mean far exceeds the census size of this population. These differences in estimates on the basis of outliers most likely result from an overcorrection when trying to exclude individuals that have been assigned to a population as a result of their geographic sampling location without genetically belonging to that population (Macbeth u. a. 2013). Including admixed individuals in the analysis of St. Lawrence Estuary resulted in $N_e$ estimates of either 989 or 185, depending on if we reduced kinship as low as possible or if we keep all individuals below first-degree relatedness. The estimation method required the exclusion of close kins; however, this is not possible in a population with such a high degree of inbreeding as the St. Lawrence Estuary population. Reducing kinship as much as possible, while retaining at least 10 samples, results in a low $N_e$ estimate of 185. An $N_e$ estimation this low is not necessarily unrealistic as previous studies have shown this population to have an $N_e$ of down to 80 (Best 2022). While it is possible that this low estimate is the result of a low sample size, a sample size of only 10 individuals also created the highest effective population size estimate for the St. Lawrence Estuary population (1,995), when we exclude the admixed individuals. It the low estimate of 185 individual can therefore not be explained simply by the small number of samples. These results demonstrate the critical importance of proper sampling when estimating contemporary $N_e$. Just the presence or absence of a small number of genetically distinct individuals from the same population can result in large differences between estimates, especially in small, strongly inbred populations like the St. Lawrence Estuary population.

**References**


Best, Brianna. 2022. „Conservation Genetics of the St. Lawrence Beluga Whale (Delphinapterus leucas)“. Bachelor Thesis, Supervisor Timothy Frasier.

COSEWIC. 2020. *COSEWIC Assessment and Status Report on the Beluga Whale, Delphinapterus Leucas, Eastern High Arctic - Baffin Bay Population, Cumberland Sound Population, Ungava Bay Population, Western Hudson Bay Population, Eastern Hudson Bay Population and James Bay Population in Canada*. Ottawa, ON: COSEWIC, Committee on the Status of Endangered Wildlife in Canada = COSEPAC, Comité sur la situation des espèces en péril au Canada.

Macbeth, Gilbert Michael, Damien Broderick, Rik C Buckworth, und Jennifer R Ovenden. 2013. „Linkage Disequilibrium Estimation of Effective Population Size with Immigrants from Divergent Populations: A Case Study on Spanish Mackerel (*Scomberomorus Commerson*)“. *G3 Genes|Genomes|Genetics* 3(4): 709–17. doi:10.1534/g3.112.005124.

Sauvé, Caroline, Pascale Caissy, Mike O Hammill, Arnaud Mosnier, und Anne P St-Pierre. 2024. „Recovery Potential Assessment for Beluga (*Delphinapterus Leucas*) Stocks in Nunavik (Northern Quebec)“.

Tinker, M T, A Mosnier, A P St-Pierre, J-F Gosselin, S Lair, R Michaud, und V Lesage. 2024. „An Integrated Population Model for St Lawrence Estuary Belugas (*Delphinapterus leucas*)“. *DFO Canadian Science Advisory Secretariat, Research Document*: iv+61 p.
